# Respiratory and Gut Microbiota in Commercial Turkey Flocks with Disparate Weight Gain Trajectories Display Differential Compositional Dynamics

**DOI:** 10.1101/2020.02.19.957092

**Authors:** Kara J.M. Taylor, John M. Ngunjiri, Michael C. Abundo, Hyesun Jang, Mohamed Elaish, Amir Ghorbani, Mahesh KC, Bonnie P. Weber, Timothy J. Johnson, Chang-Won Lee

## Abstract

Host-associated communities of bacteria (microbiota) substantially contribute to the overall poultry health and performance. Gut microbiota are known to play roles in resistance to pathogen infection and optimal weight gain in turkey flocks. However, knowledge of turkey respiratory microbiota and its link to gut microbiota is lacking. This study presents a 16S rRNA gene-based census of the turkey respiratory microbiota (nasal cavity and trachea) alongside gut microbiota (cecum and ileum) in two identical commercial Hybrid Converter turkey flocks raised in parallel under typical field commercial conditions. The flocks were housed in adjacent barns during the brood stage and in geographically separated farms during the grow-out stage. Several bacterial taxa that were acquired in the respiratory tract (RT) at the beginning of the brood stage persisted throughout the flock cycle, primarily *Staphylococcus*. Late-emerging predominant taxa in RT included *Deinococcus* and *Corynebacterium*. Tracheal and nasal microbiota of turkeys were identifiably distinct from one another and from gut microbiota. Nevertheless, gut and RT microbiota changed in parallel over time and appeared to share many taxa. During the brood stage, the two flocks generally acquired similar gut and RT microbiota, and their average body weights were comparable. Separating the flocks during the grow-out stage resulted in divergent microbial profiles and body weight gain trajectories. Lower weight gain corresponded with emergence of *Deinococcus* and *Ornithobacterium* in RT, and *Fusobacterium* and *Parasutterella* in gut. This study provides an overview of turkey microbiota under field conditions and suggests several hypotheses concerning the respiratory microbiome.

**IMPORTANCE:** Turkey meat is an important source of animal protein, and the industry around its production contributes significantly to the agricultural economy. The nonpathogenic symbionts present in the gut of turkeys are known to impact bird health and flock performance. However, the respiratory microbiota in turkeys are entirely unexplored. This study has elucidated the microbiota of respiratory tracts of turkeys from two commercial flocks raised in parallel throughout a normal flock cycle. Further, the study suggests that bacteria originating in the gut or in poultry house environments may influence respiratory communities and consequently induce poor performance, either directly or indirectly. Future attempts to develop microbiome-based interventions for turkey health should delimit the contributions of respiratory microbiota and aim to limit disturbances to those communities.

## INTRODUCTION

Bacterial communities (microbiota) associated with poultry have great influences on bird development (1, 2), health (3–5), and production performance (6, 7). A wealth of information on microbial density and composition in the gastrointestinal tracts of broiler chickens, layer chickens, and turkeys has been generated in the last decade (3, 4, 8–10). Certain bacterial species in the turkey gut are associated with adverse effects such as increased susceptibility to pathogens (11–13), inefficient feed conversion (1), and suboptimal market weights (6, 14), while other species are correlated with improved health and enhanced performance metrics, including weight gain (1, 6, 15, 16).

Microbiota composition in the turkey gut is dependent on a number of intrinsic and extrinsic factors. Intrinsic factors such as bird age and the site within the gut (cecum, ileum, duodenum, jejunum, ileum, etc.) are good predictors of the diversity and composition of communities in commercial turkeys (6, 17, 18). Gradual parallel composition changes occur in different gut sites as the bird ages (6, 17, 18), reflecting anatomical differences between sites and age-dependent physiological changes. Possible extrinsic factors are farm location (19), flock rearing and health management practices (15, 20, 21), including diet, housing conditions (22), litter reuse (10, 23, 24), and vaccination and antibiotic use (25). These factors do cause microbiota composition changes, but the effects of those changes on production performance metrics are variable and thus difficult to predict. In particular, we have shown that the gut microbial composition of multiple chicken flocks of the same breed and raised adjacently may differentiate from one another with age (26), potentially leading to a performance gap between flocks. Such detailed information is lacking for turkey flocks.

A thorough understanding of the gut and respiratory microbiota is critical to decipher the potential roles of microbiota in pathogen resistance in addition to production performance. Our previous work in chickens demonstrated that the microbiota of the respiratory system overlap substantially with gastrointestinal microbiota (26, 27). We observed that numerous taxa reside simultaneously in the respiratory and gastrointestinal tracts and shift in parallel over time (26, 27). It is vitally important to establish whether these dynamics are true in turkeys as well, as they may indicate how abrupt changes to flock management disrupt both systems.

Currently, there is no exposition of turkey respiratory microbiota that would permit comparison with gut microbiota. Previous studies of turkey respiratory bacteria have focused on specific pathogens, such as *Mycoplasma* (28, 29) and *Ornithobacterium rhinotracheale* (30) and have not considered non-pathogenic microbial communities residing therein. In other poultry, different respiratory sites are known to house distinct bacterial communities that develop with age (26, 27, 31, 32), but the contribution of these communities to poultry health is largely unknown. Of particular interest are the microbiota of the upper respiratory system (the nasal cavity and the trachea), as these sites are among the first points of contact for airborne bacteria, including aerosolized fecal bacteria (33). Understanding how upper respiratory microbiota develops alongside lower gastrointestinal microbiota is critical to the development of effective interventions against transmission of performance-inhibiting pathogens and to improve poultry health and productivity.

In the current study, we provide respiratory and gastrointestinal microbiota observed in two identical commercial turkey flocks from a vertically integrated turkey production system. The birds were raised in adjacent barns throughout the brood farm stage but were geographically separated for the grow-out/finisher farm stage. This management setup allowed parallel observation of weight gain and microbiota both before and after separation and transfer to new locations.

## RESULTS

### Health status and performance of the flocks

Both flocks were vaccinated against HEV and *Salmonella* (between 3 and 4 weeks of age, WOA) before transition to the grow-out farms. During the grow-out/finish stage, while both flocks were booster-vaccinated against *Salmonella*, flock F1G was also administered with *Pasteurella multocida* vaccine. At 1 WOA, the flocks had high levels of antibodies against HEV, NDV, REO, and BA that are presumed to have been passively transferred from the breeder hens to the poults through the egg yolk (Fig. 1B). The maternally-derived antibodies subsequently waned to low or undetectable levels unless the flock was vaccinated (HEV) or became infected (REO). Antibody response to HEV vaccination was minimal between 5 and 8 WOA but robust during 12 and 16 WOA. The presence of increasing titers of anti-REO antibodies from the 3^rd^ WOA onward suggests the flock was infected with reovirus of unknown pathogenicity (Fig. 1B), as has been reported in other commercial flocks (34).

**Fig. 1.**
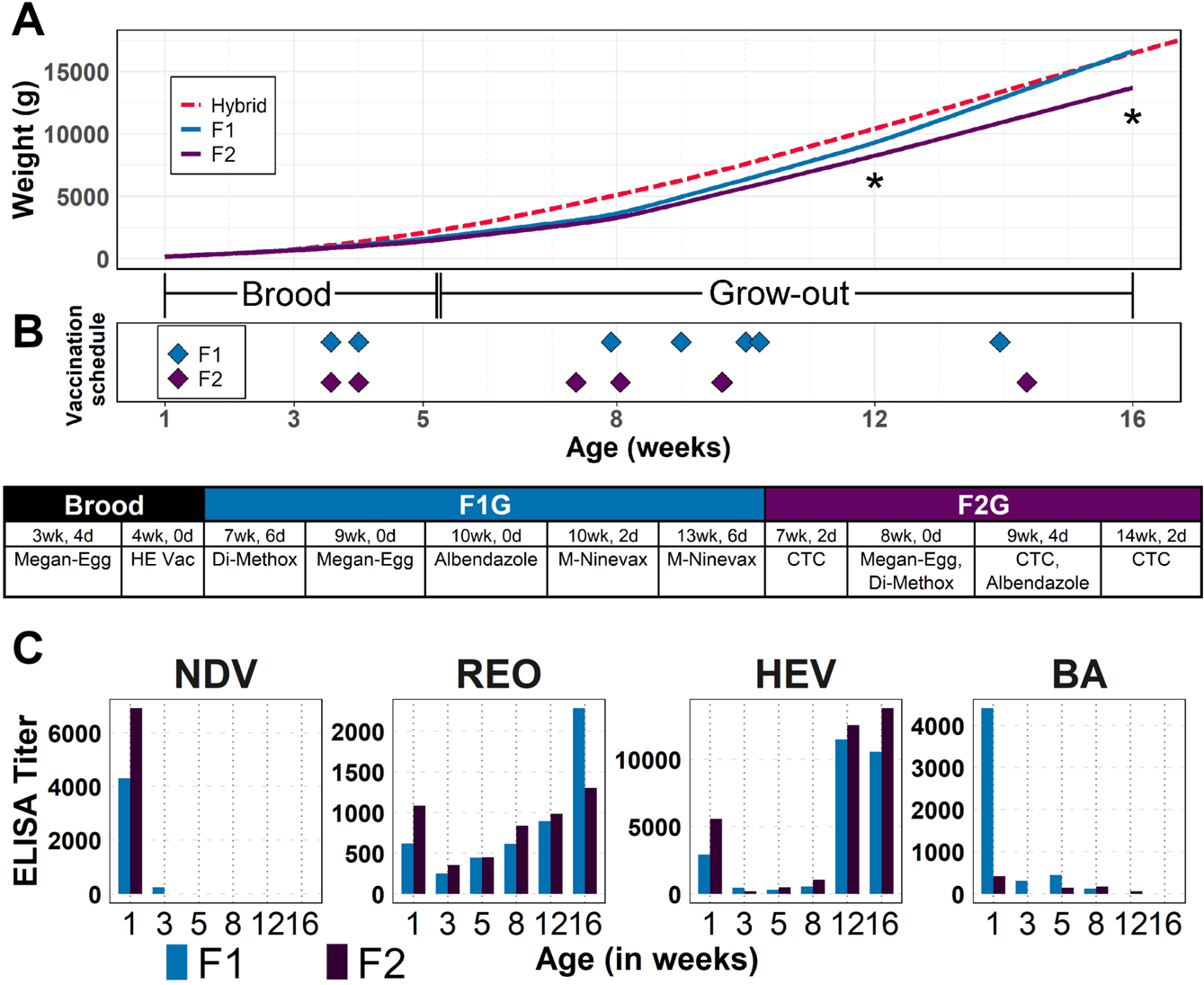
Flock metadata: Body weights, vaccination timeline, and immune status. Flocks F1 and F2 were raised by the same management during the brood stage (weeks 1-5) and separated to two different farms at the grow-out stage (after week 5). **(A)** Average weights for each flock (F1 in blue, F2 in purple) compared to expected weights for the Hybrid Converter breed (red). Asterisks indicate ages at which bird weights were significantly different between F1 and F2 (p<0.001). **(B)** Timing of vaccinations during the sampling period (top) and exact ages and compounds used (bottom). **(C)** Antibody titers for Newcastle Disease virus (NDV), reovirus (REO), hemorrhagic enteritis virus (HEV), and *Bordetella avium* (BA). Bird ages-at-sampling are given on the x-axes of (B) and (C).

We did not determine the immune responses to *Salmonella* and *Pasteurella multocida* vaccinations, and no ASVs were identified as *Salmonella* or *Pasteurella* before or after vaccination. Both flocks were serologically negative for *Mycoplasma gallisepticum*, *M. meleagridis*, and *M. synoviae,* well known pathogens for turkeys (35). In addition to vaccination, the flocks were treated with albendazole (antihelminthic) and sulfadimethoxine (coccidiostat-antibacterial) as part of routine management protocol.

Production performance was tracked by taking body weights of the sampled birds. Starting from 5 WOA onward, the average body weight of flock F2 were below the genetic potential of the Hybrid Converter (36) and remained so until the end of study timeline (Fig. 1A). A similar trend of lower relative body weight gain was observed in flock F1G until 8 WOA when the body weights started to gain rapidly and reached to genetic potential by 16 WOA. Notably, the average body weight of flock F2G was significantly lighter than that of flock F1G at 12 and 16 WOA (Fig. 1A).

### Census of predominant bacterial taxa distinguishes between the lower intestinal tract (LIT) and upper respiratory tract (URT) environments

In the cecum, the dominant phyla were Firmicutes and Bacteroidetes, largely represented by the orders Clostridiales and Bacteroidales, respectively (Fig. S1). The most prominent genus was *Bacteroides*, although *Lactobacillus* was also common during the brood stage (Fig. S2). In the ileum, the predominant phylum was Firmicutes, with the dominant orders being Clostridiales and Lactobacillales, and the most prominent genus being *Lactobacillus*. ASVs classified as *Candidatus* division *Arthromitus* were abundant (>5% relative abundance) at 1WOA, but rapidly declined to <1% relative abundance by 12 WOA (Fig. S3A).

Predominant taxa in the respiratory tract of turkeys were classified as *Deinococcus*, *Corynebacterium*, and *Staphylococcus* (see Fig. S8-S11). Potentially pathogenic ASVs from the genera *Mycoplasma* and *Ornithobacterium* were abundant in both respiratory sites (Fig. S3B), even though serology for well-known *Mycoplasma* species was negative. At the phylum level, Firmicutes and Actinobacteria predominated in the nasal cavity. The distribution of tracheal phyla through time was not as consistent as in the other three body sites (Fig. S1); however, there was a much higher prevalence of Proteobacteria than in other sites.

In sum, each body site was characterized by the predominance of a particular genus at all sampling times: *Bacteroides* in cecum, *Lactobacillus* in ileum, and *Staphylococcus* in the nasal cavity, but no genus was consistently predominant in trachea (Fig. S2).

### Microbial communities mainly assemble according to body sites and body systems

The four body sites sampled were measurably different according to several exploratory indices. The total number of ASVs observed was higher in LIT (1434 in cecum and 1145 in ileum) than in URT (717 in nasal cavity and 689 in trachea) (Fig. S5). However, the average bacterial species richness in each bird was higher in cecum (209 ASVs) and nasal cavity (105 ASVs), and lower in ileum (89 ASVs) and trachea (66 ASVs) (Fig. S5).

In terms of overall composition of bacterial communities among the sampled birds, the four body sites are distinct from one another, and the composition of each site changes in various ways as the birds age. Based on principal coordinates analysis (PCO) of unweighted UniFrac sample distances, samples clustered according to sampling age (PERMANOVA, p<0.001) and body site (PERMANOVA, p<0.001). The distinction between groups was visible in Fig. 2A, with 44.5% of total variance explained (PC1=25.7%, PC2=10.8%, PC3=8.0%). Qualitatively, PC1 separated samples into URT and cecum, with ileum samples distributed along the range (Fig. 2A), and PC2 stratified samples by age, regardless of body site (Fig. S4A).

**Fig. 2.**
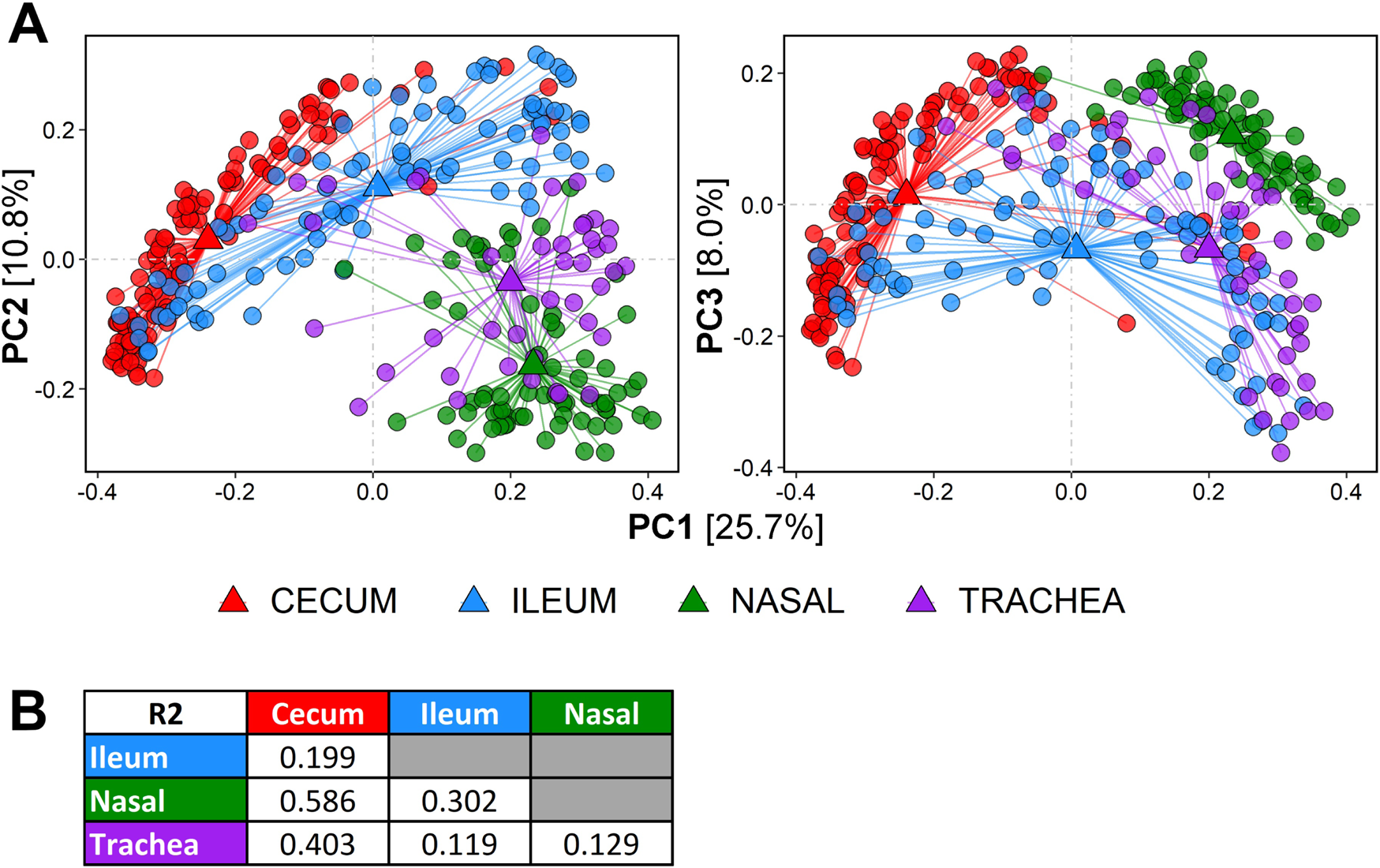
Comparison of bacterial community composition in URT and LIT sites. **(A)** Principal coordinates plot of sample distribution according to unweighted UniFrac distances. Axes displayed are PC2 (left plots) and PC3 (right plots) with reference to PC1. Top row shows samples colored according to body site, bottom row shows samples colored according to age-at-sampling. Values in brackets beside axes indicate the percentage of sample variance explained by that axis. Total variance explained by PC1, PC2 and PC3 is 44.5%. **(B)** R^2^ values from pairwise PERMANOVA comparison. The R^2^ quantifies how well the grouping variable describes the distances between group centroids (triangles). Larger R^2^ values indicate greater distinction between groups.

Quantitative distinctions between sample groups are specified by PERMANOVA as the amount of variance explained (R^2^) by the distance between the centroids of pairs of groups. All body sites were significantly distant from one another (PERMANOVA, p<0.001 for all comparisons) (Fig. 2A), but the R^2^ values were variable (Fig. 2B). Centroid distances were smaller within LIT (R^2^=0.199) and within URT (R^2^=0.129). Cecum and URT sites were the most distant (R^2^ with nasal=0.586; R^2^ with trachea=0.403); followed by ileum and nasal (R^2^=0.302). Trachea and ileum were the least distant (R2=0.119) (Fig. 2B).

Age had a significant effect on sample grouping when all four body sites were ordinated together (Fig. S4A) and when each body site was ordinated independently (Fig. 3; PERMANOVA, p<0.001 for all sites). Total variance explained by the first two axes is 44.8% in cecum, 38.0% in ileum, 31.2% in nasal cavity, and 37.6% in trachea. Sample separation between brood (1-5 WOA) and grow-out (8-16 WOA) stages is visible in all four body sites (Fig. 3).

**Fig. 3.**
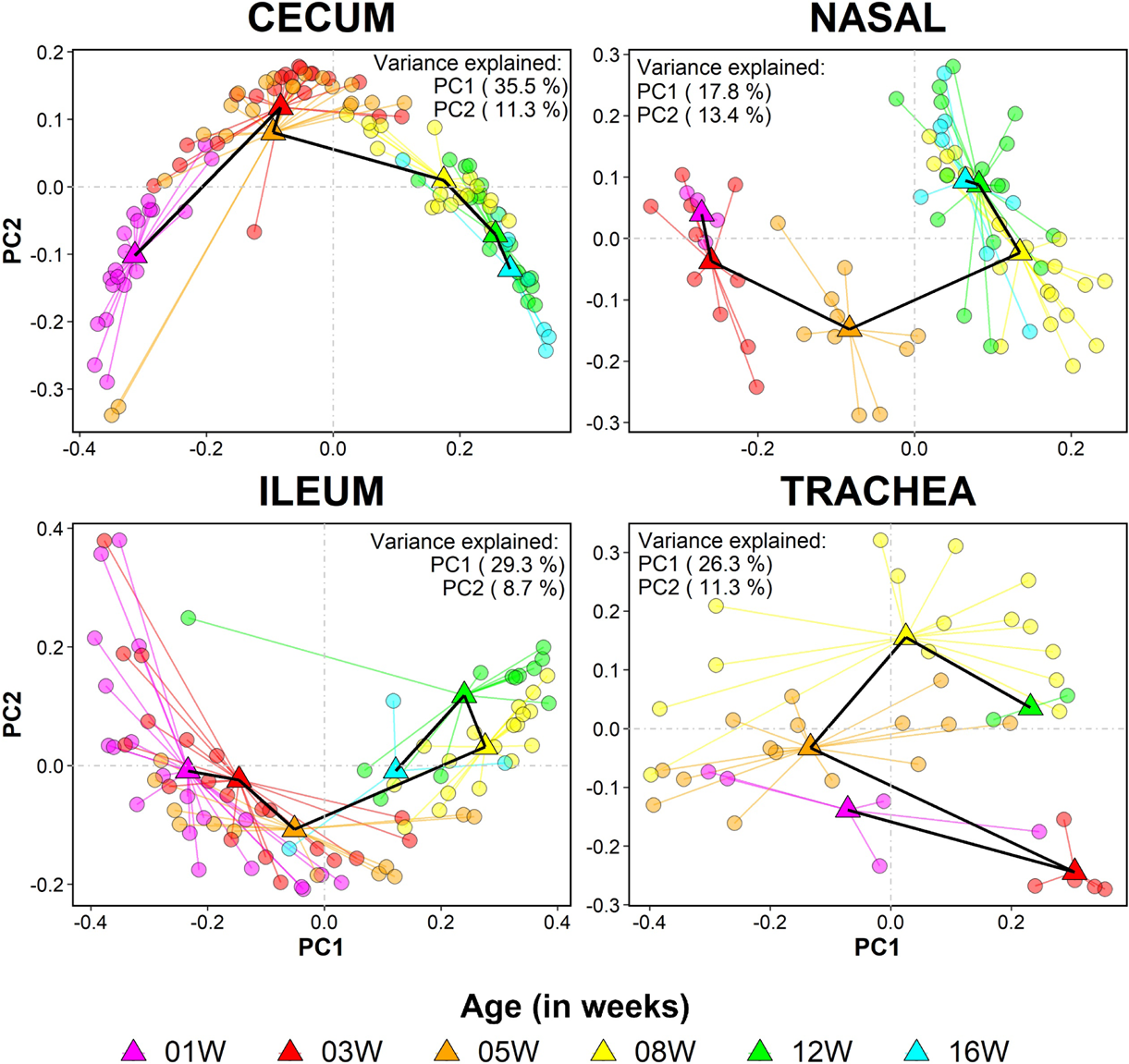
Community composition of samples from each body site changes gradually with host age. Principal coordinates plot of sample distribution given according to unweighted UniFrac distances. Samples (circles) are colored according to age-at-sampling: 1WOA (pink), 3WOA (red), 5WOA (orange), 8WOA (yellow), 12WOA (green), and 16WOA (blue). The black line connects the centroids of consecutive age groups. Only the first two axes are given for each plot. Variance explained by each axis is given in the corner of each plot.

### LIT and URT communities do not exist in isolation from each other

Despite obvious differences in bacterial community composition and species richness between body sites (Fig. 2, see text above), several taxa were shared among them. Of the 1,874 ASVs observed in this study, 285 were shared across all four body sites, 506 were shared only between LIT sites (cecum and ileum), and 133 were only shared between URT sites (trachea and nasal cavity) (Fig. S5). The consensus relative abundances of shared and unique ASVs vary according to body site. Hereafter, we refer to the total potential relative abundance of ASVs present in a body site as “sample space” or “percent sample space” (the maximum of which is 100%) as a means of describing site occupancy without differentiating transient species from long-term inhabitants. The sample space occupied by an ASV is presented as a mean relative abundance for that ASV averaged across all samples of the same age group, body site, and flock (unless flock is not indicated in the figure; e.g. Fig. 4). The sample space occupied by shared ASVs is summarized in Fig. 4A and in greater detail in Fig. S5. Irrespective of age, ASVs that were shared amongst all body sites occupied more sample space in LIT than URT (t-test, p<0.01; Fig. 4B). In general, as the amount of sample space occupied by the ubiquitous ASVs decreased with age, there was complementary increase in the sample space occupied by ASVs unique to LIT sites or URT sites only (Fig. 4A, Fig. S5).

**Fig. 4.**
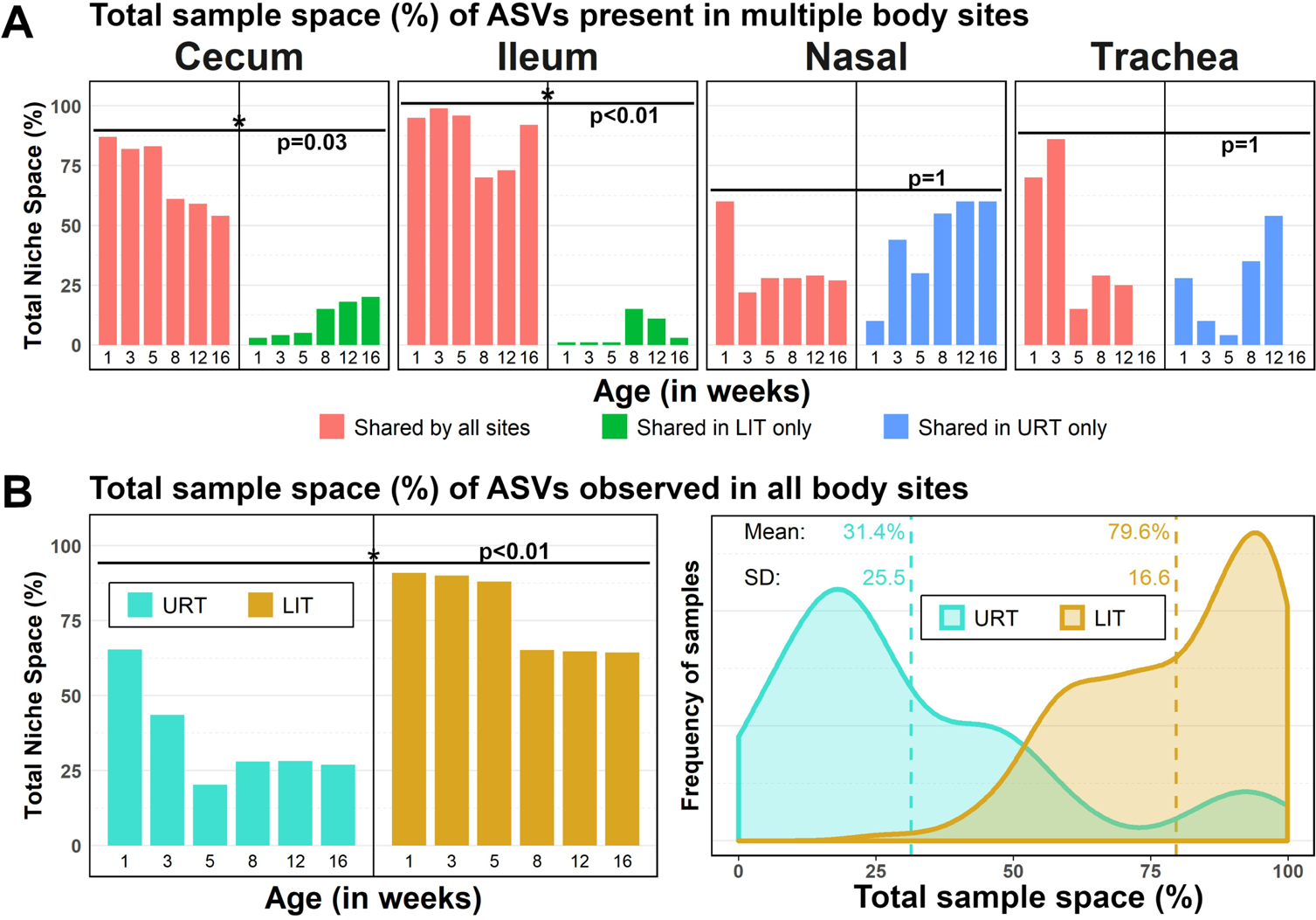
The total sample space (per age group) occupied by ASVs that are shared by all sites (red; 285/1874 ASVs), shared by sites in the URT only (blue; 133/1874 ASVs), and shared by sites in the LIT only (green; 506/1874 ASVs). Percentages given are the mean total relative abundance of shared ASVs in a body site at a particular sampling age. Paired t-tests were performed between the mean total abundance of ASV groups per age in a pairwise fashion with Bonferroni correction for multiple comparisons: **(A)** between ASV groups in a body site, **(B, left)** between URT and LIT. Asterisks indicate significant differences between ASV groups (p<0.05). **(B, right)** The distribution of sample values for total sample space occupied by ubiquitous ASVs is also shown to demonstrate the distinction in URT and LIT occupancy by these ASVs at the sample level. A Venn diagram of ASV distribution among samples is given in **Fig. S5**.

### Flock barn and grow-out stage separation had minimal influence on alpha diversity, except in the nasal cavity

The flocks were housed in two proximally located barns within the same farm and geographical location during the brood stage, designated F1B & F2B. The birds of these brood barns were destined for different (geographically distant) grow-out farms, designated F1G & F2G, respectively. Age-independent examination of species richness (measured as the number of ASVs observed) and Pielou’s evenness did not significantly differentiate the two flocks in any body site. However, species richness distribution indicated significant differences between the brood and grow-out stages (t-test: p<0.05), but not between flocks within each stage (t-test: p>0.1). This pattern was consistent in all four body sites. Linear regression of species richness and evenness against week-of-age-at-sampling indicated mostly minor differences between flock groups (Table 1 and Fig. S6). Species richness in individual birds tended to increase with age (*b*>0, where *b* is the slope). Although richness correlated well with age-at-sampling for LIT samples (cecum R^2^>0.6; ileum R^2^>0.2), it did not correlate consistently well with URT samples. Specifically, species richness in the nasal cavities of F2 birds correlated positively with age-at-sampling (*b*=6.23, R^2^=0.53), but fluctuated frequently in F1 birds (*b*=-0.24, R^2^<0.01). The slopes of each correlation were not statistically different between F1 and F2 in cecum, ileum, or trachea, but were significantly different in nasal cavity (t-test, p<0.001; Table 1 and Fig. S6). Bacterial community evenness in individual birds tended to increase with increasing sampling age, but the correlations were generally not as strong as with richness. The changes in evenness over time in the ileum and trachea were not significantly different between F1 and F2 birds. However, significant differences were seen in the cecum and nasal cavity (Table 1 and Fig. S6).

**Table 1.** Coefficients* for linear regression models between richness and evenness indices and bird <u>sampling age</u>

|  | Body Site | Flock | *b* | *A* | *R*<sup>2</sup> | *df* | *v* | *T* | *p* |
| --- | --- | --- | --- | --- | --- | --- | --- | --- | --- |
| Richness | Cecum | F1 | 20.7 | 79.6 | 0.863 | 51 | 1.33 | 1.07 | 0.224 |
|  |  | F2 | 16.4 | 103 | 0.663 | 51 | 2.68 |  |  |
|  | Ileum | F1 | 13.6 | 20.2 | 0.549 | 39 | 3.90 | 0.70 | 0.310 |
|  |  | F2 | 8.32 | 37.2 | 0.26 | 44 | 4.48 |  |  |
|  | Nasal | F1 | -0.24 | 105 | 0.00159 | 33 | 1.10 | 9.97 | 1.73E-14 |
|  |  | F2 | 6.23 | 56 | 0.53 | 33 | 1.04 |  |  |
|  | Trachea | F1 | 4.3 | 59 | 0.0561 | 21 | 14.8 | 1.09 | 0.217 |
|  |  | F2 | 3.77 | 25.6 | 0.0851 | 19 | 8.04 |  |  |
| Evenness | Cecum | F1 | 0.00475 | 0.698 | 0.196 | 51 | 1.81E-06 | 3.12 | 0.004 |
|  |  | F2 | 0.00916 | 0.647 | 0.227 | 51 | 5.60E-06 |  |  |
|  | Ileum | F1 | 0.0216 | 0.482 | 0.337 | 39 | 2.35E-05 | 0.34 | 0.375 |
|  |  | F2 | 0.00966 | 0.5 | 0.077 | 44 | 2.54E-05 |  |  |
|  | Nasal | F1 | -0.00268 | 0.645 | 0.0351 | 33 | 5.98E-06 | 9.49 | 1.20E-13 |
|  |  | F2 | 0.0101 | 0.573 | 0.5 | 33 | 3.09E-06 |  |  |
|  | Trachea | F1 | 0.0116 | 0.425 | 0.0163 | 21 | 3.87E-04 | 0.65 | 0.319 |
|  |  | F2 | 0.00629 | 0.371 | 0.00723 | 19 | 2.86E-04 |  |  |

\* Coefficients given are slope (*b*), y-intercept (*A*), coefficient of determination (*R*<sup>2</sup>). Also given are the degrees of freedom for the group (*df* = *n* - 1), variance around the slope (*v*; see **Function 1**), and an associated value for Student's *T* comparing the two slopes (*T*; see **Function 2**), from which a p-value is derived (*p*).

### Bacterial community compositions were differentiated after the flocks transitioned to the grow-out barns

Like species richness and evenness, the differences in bacterial species composition between birds (as measured by unweighted UniFrac distances) visibly changed with age in all body sites (see Fig. 3). However, bacterial communities of each body site in the two flocks were compositionally more similar at the brood stage than at the grow-out stage in the cecum, ileum, and nasal cavity (PERMANOVA, p<0.05, Table 2 and Fig. S7). In contrast, tracheal composition was more differentiated at the brood stage than in the grow-out stage. The composition of each body site was further examined by focusing on the dynamics of several predominant ASVs across age-groups. ASVs were considered “predominant” if they occurred in 50% of birds within an age-group, were retained from emergence until 16 WOA, and achieved a relative abundance of >3.5% in at least one age-group. The relative abundances of constituent bacteria that were predominant in both groups during the brood stage were differentially decreased by the acquisition of new ASVs at the start of the grow-out stage (i.e. post-transfer to new farms) (Fig. S8-S11).

**Table 2.** Statistical comparison of unweighted UniFrac distances between flocks and farm stages

| **Body Site** | **Between Brood<sup>†</sup>**<br>(F1B vs F2B) | **Between Grow-out<sup>†</sup>**<br>(F1G vs F2G) | **Brood (F1B+F2B) vs.**<br>**Grow-out (F1G + F2G)** | **Between Flock<sup>†</sup>**<br>(Brood + Grow-out) |
| --- | --- | --- | --- | --- |
| Cecum | 0.006<sup>#</sup> | 0.276* | 0.412* | 0.049 |
| Ileum | 0.026 | 0.148* | 0.403* | 0.023 |
| Nasal | 0.020 | 0.134* | 0.327* | 0.013 |
| Trachea | 0.195* | 0.157 | 0.084* | 0.015 |

Generally speaking, a greater number of bacteria gained predominance at the grow-out stage in URT (particularly in trachea) than LIT, regardless of flock group. Most predominant genera in the nasal cavity persisted from 1WOA (F1+F2: *Brevibacterium*, *Corynebacterium*, *Lactobacillus*, *Rothia*, *Staphylococcus*; F1: *Kocuria; F2: Macrococcus*). From 8 WOA, F2 acquired *Deinococcus*.

In general, very few predominant taxa persisted long-term in trachea compared to the nasal cavity. Members of Burkholderiaceae, *Escherichia*, *Lactobacillus, Staphylococcus* (F1+F2), *Bifidobacterium*, and *Streptococcus* (F1) all appeared in the trachea prior to farm transfer. *Corynebacterium* (F1+F2), *Macrococcus* (F1), *Brevibacterium*, *Deinococcus*, and *Rothia* (F2) emerged and persisted in trachea from 8 WOA (after the transfer). (Fig. S8-S11). Strikingly, *Deinococcus* only became predominant in the URT of F2G birds (Fig. S10-S11). Differences in *Deinococcus* abundance between groups were tested with Dunn’s Kruskal-Wallis rank sum test and Holm’s correction for multiple comparisons. The abundance of *Deinococcus* in nasal and tracheal samples was not different between F1B, F2B, and F1G (p>0.2). Conversely, *Deinococcus* abundance was significantly higher in F2G compared to the other groups in both nasal and tracheal samples (p<0.009).

In the LIT, genera that tended to emerge at 1WOA and persist through 16WOA were *Bacteroides*, *Lactobacillus*, *Ruminococcus*, *Bifidobacterium*, and *Escherichia*. Genera that emerged as predominant in LIT after farm transfer (8-16WOA) were *Phascolarctobacterium*, *Sutterella*, *Alistipes*, and *Romboutsia* (Fig. S8-S11). Although several genera predominated simultaneously between flocks, the individual ASVs representing those genera in LIT and URT were rarely shared between the flocks.

### New acquisition of unique taxa coincides with the transfer to new farms

Differences between groups were minimal during the brood stage but increased after transfer to grow-out barns: ASVs shared between the four flock groups (483/1874) occupied nearly 100% of available sample space during the brood stage, but that percentage was reduced in both flocks after the move (Fig. 5A). New ASVs were acquired in both flocks after the farm transfer, 489 of which were common to both flocks, 273 of which were unique to F1G, and 161 of which were unique to F2G (Fig. 5B; see Fig. S12 for the most abundant ASVs in each flock and Data Sheet S1 for complete ASV list). The most abundant ASVs unique to F1G were classified as *Fusobacterium*, *Muribaculaceae*, *Alistipes*, *Mucispirillum*, *Methanobrevibacter*, *Staphylococcus*, *Macrococcus*, *Rothia*, *Lactobacillus avarius*, *Peptococcus*, *Lactobacillus*, *Mycoplasma*, and *Eikenella* (Fig. 5C). The most abundant ASVs unique to F2G were classified as *Parasutterella*, *Faecalibacterium*, *Caproiciproducens*, *Ruminococcaceae*, *Mollicutes*, *Olsenella*, *Clostridium butyricum*, *Escherichia-Shigella*, *Deinococcus*, *Deinococcus piscis*, *Rothia*, *Chryseobacterium*, *Acidipropionibacterium*, and *Paracoccus* (Fig. 5C).

**Fig. 5.**
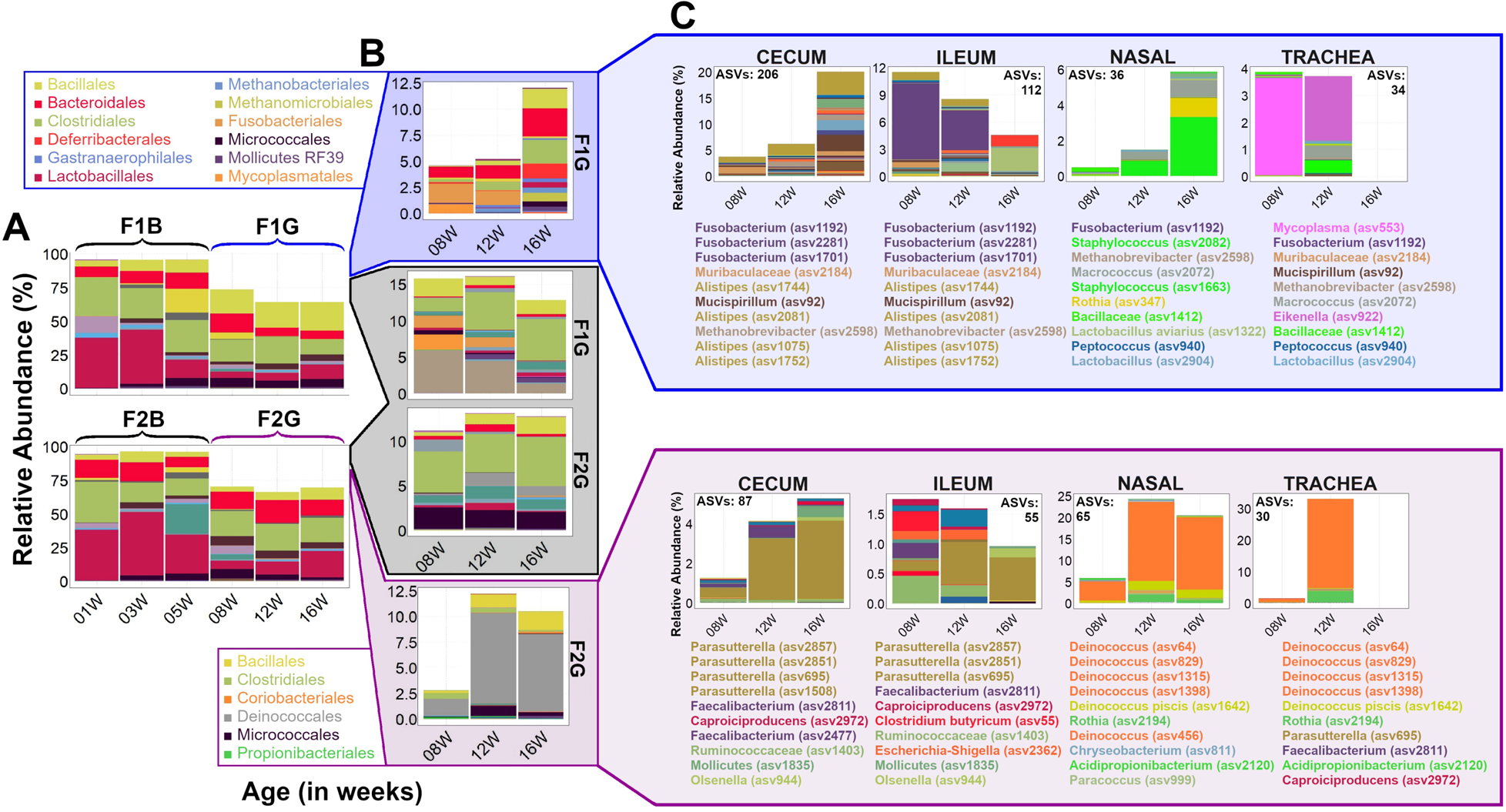
Dynamics of unique bacterial taxa that are acquired by each flock independently during the grow-out period. **(A)** The total sample space occupied by ASVs that were common to both flocks during both the brood and grow-out stages. Colors represent different taxonomic orders. Note that the sample space occupied is close to 100% in the brood stage but is depressed in the grow-out stage. **(B)** The majority of remaining sample space is occupied by ASVs either shared amongst the grow-out flocks (black box), or unique to each (blue and purple boxes). Colors represent different taxonomic orders, the most abundant of which are given in the adjacent legends. **(C)** The total relative abundance of ASVs unique to each flock, distributed between the 4 body sites. Legends below each bar chart name the top 10 most abundant of these ASVs within each body site. Colors represent the lowest taxonomic level assigned to an ASV by the SILVA classifier.

### Potentially pathogenic bacteria identified in both flocks

We identified two ASVs that closely matched *M. gallisepticum*, *M. tullyi*, and *M. imitans* when referenced against the NCBI Blast 16S rRNA gene database (see Fig. S3 and S12). It is likely these two ASVs represent *M. tullyi* (37) or *M. imitans* (38, 39) since both flocks were serologically negative for *M. gallisepticum*. Both *Mycoplasma* ASVs were present only at 8 WOA. Additionally, the common respiratory pathogen *Ornithobacterium rhinotracheale* was observed as a single ASV that matched the Blast reference for *O. rhinotracheale* with 100% identity in both flocks. *O. rhinotracheale* occupied 68.9% of tracheal sample space in F2 at 5 WOA, dropping to 22.1% at 8 WOA, while occupying 7.2% of sample space in F1 (8 WOA) (see Fig. S3).

## DISCUSSION

Information on the turkey respiratory tract microbiota is currently limited, as no culture-independent surveys have been conducted previously to the best of our knowledge. This study provides an extensive census of bacterial residents in both the URT (trachea and nasal cavity) of turkeys in conjunction with the LIT (ileum and cecum), in the course of a typical commercial flock cycle. Bacterial community compositions among the body sites formed a gradient, with highly structured communities (cecum and nasal cavity) at the extremes and less structured communities (ileum and trachea) in between (Fig. 2). This gradient closely recapitulates our recent observations in layer and broiler chickens (26, 27). Additionally, there was substantial sharing of bacterial taxa between the gut and RT (Fig. 4 and S5). The sample space occupied by these taxa was significantly higher in the gut than in RT (Fig. 4), suggesting that several of these taxa may colonize the gut and migrate to the RT as aerosolized fecal matter. What is yet unclear is how many of those migrating taxa are ephemeral occupants versus long-term inhabitants of respiratory mucosal layers. The baseline data generated in this study will guide future controlled experiments designed to uncover causal relationships between respiratory microbiota, gut microbiota, and turkey flock performance.

The presence or absence of certain respiratory pathogens coincided with the emergence of below-average weight during the late brood and early grow-out period (5-8 WOA). For instance, the respiratory pathogen *O. rhinotracheale* (1 ASV) (35, 40, 41) occupied >60% of tracheal sample space in F2 at 5 WOA (Fig. S3B), a trend that diminished with subsequent samplings, but likely influenced the stall in weight gain recovery in that flock (42). *Mycoplasma* could be significant in the context of below-average rates of weight gain observed in flocks (29). Although serological evidence precluded the presence of antibodies to MG, MM, and MS, flock F2 began receiving precautionary treatment with CTC at 7 WOA. This treatment may account for the low relative abundance of *Mycoplasma* ASVs in F2 relative to F1 at 8 WOA (Fig. S3B), yet the abundance of Mycoplasma in each flock inversely corresponded to disparities in average weight (Fig. 1). It is possible that the effects of CTC on composition are compounded on the effects of the farm move, resulting in a prolonged period of microbiota disruption.

After 8 WOA, the flocks began to differentiate in their trajectories of weight gain (Fig. 1A) and at the same time gained several bacterial taxa unique to each flock (Fig. 5). In F1G the trend of weight recovery visible from 12 WOA coincided with *Staphylococcus* increase in the nasal cavity. The novel *Staphylococcus* ASVs (Fig. 5) in the nasal cavity of F1G joined a large extant population of these bacteria (Fig. S10). Although several *Staphylococcus* species are known pathogens (43), abundance of *Staphylococcus* increased as bird weight improved in F1 and remained suppressed in F2 (where bird weight did not improve). This suggests that some *Staphylococcus* species may behave commensally, by potentially preventing colonization of the mucosal layer by naïve environmental bacteria.

The lack of weight recovery in F2G also coincided with the emergence and persistence of numerous *Deinococcus* ASVs in the respiratory tract (Fig. 1A and 5). *Deinococcus* has been previously identified as a common extraction kit reagent contaminant, particularly in low biomass samples (44). Given that RT wash samples are typically low biomass (compared to gut content samples) (44), the prevalence of this genus in the RT samples of F2G was initially cause for concern. However, birds of the same age group from both flocks were necropsied at the same time, and the processed samples were extracted utilizing the same aliquots of reagents. Under these conditions, contaminants are likely to be well distributed among samples processed together, and this was not observed. The significantly greater abundance of *Deinococcus* in F2G suggests a contaminant in the housing itself that may be inhabiting mucosal surfaces in the URT. *Deinococcus* species have been previously isolated in a number of poultry environments, including rearing houses and carcass processing facilities, even after decontamination (45, 46).

This genus has also been shown to carry the tetracycline-efflux transporter gene required for resistance to CTC (47), which suggests that it may have expanded as CTC-susceptible taxa declined in this flock (Fig. 1). Its effects on poultry health have not been explored in depth, although one study reported that supplementing layer hen feed with dried-fermented powder of *Deinococcus* improved feed conversion and egg yolk color (Li et al., 2018). Their pervasiveness in the respiratory system may ultimately be innocuous, but any causal relationship with weight suppression and possible mechanisms of disruption can only be elucidated under controlled conditions.

The associations of gut microbiota to poultry performance metrics are well studied (6, 48–50). Changes in the abundance of certain taxa after a disturbance (in this case, transfer to new barns or farms) are often associated with performance issues and pathogen infections (3, 5, 22, 25, 51). For example, the rapid decline of *Candidatus* division *Arthromitus* in ilea of both flocks during the brood stage (Fig. S3) corresponded qualitatively with emergence and persistence of below average weights (Fig. 1). A causal relationship between *Candidatus* extinction and poor weight gain in young poults is likely since this taxon was previously found to be dominant in turkey flocks characterized as “Heavy” (6, 16, 25).

As in the trachea and nasal cavity, there were several gut bacteria that were uniquely acquired by each flock at the grow-out farms. *Fusobacterium* ASVs were abundant in flock F1 ilea at 8 WOA (when weight gain was most suppressed) and gradually diminished to very low levels by 16 WOA (when average bird body weight reached the expected weight) (Fig. 1 and 5). F2G experienced noticeable increases of *Parasutterella* ASVs in cecum and ileum as body weights failed to recover between 8 and 16 WOA. Both *Fusobacterium* and *Parasutterella* spp. have been identified in the poultry gut (48, 52–54) but with no overt links made to poor performance. However, both genera have been associated with pathogenic effects in humans and other species (55–58). It might be worthwhile to examine these genera more closely with reference to potential correlations with poultry weight or other metrics of performance.

Beyond the potential associations with trajectories of weight gain, important insights into turkey microbiota beta diversity were indicated. Firstly, this study corroborates previous observations that turkey age is a discernible driver of compositional change in LIT sites (Fig. 3) (6, 17, 18). Age is also an important driver of respiratory microbiota, but more so in the nasal cavity than in trachea. Secondly, although communities of all body sites shifted significantly with the farm move (Table 2; Fig. S7), differentiation between the flocks was only observed in the LIT and nasal cavity. This indicates that the avian LIT and nasal cavity environments are more hospitable to long-term inhabitants than the trachea (59). Community composition in the avian trachea may be mostly ephemeral and turn over species more frequently, as observed in the human respiratory tract (60–62).

In contrast with beta diversity, alpha diversity trends with age were notably different between the LIT and URT. In cecum and ileum, species richness increased as linear functions of age that did not significantly differ between flocks (Table 1). Bacterial alpha diversity change with age in gut sites has been shown previously in poultry and humans (63–65) but evidence of similar trends in poultry respiratory sites is lacking. In this study, neither the nasal cavity nor the trachea demonstrated a consistent linear relationship between alpha diversity (species richness and evenness) and turkey age (Table 1). After the farm transfer, diversity in the nasal cavity remained constant in F1 but suddenly increased in F2, possibly as a result of high diversity in the barn environment serving as the source population (23, 24, 66). Tracheal diversity was not as systematically affected by the farm transfer in either flock, although several nasal bacteria were also observed in the trachea. At this time, it is unclear whether one of the two sites acquired bacteria first or whether they acquired bacteria simultaneously and share continuously. Further study of microbial exchanges between respiratory sites is fundamental for the development of intervention strategies towards reducing pathogen load and recuperating poor performance.

It should be noted that while age-dependent microbial dynamics in different body sites are the conventional focus of poultry microbiome research and are significant drivers of compositional variation (6, 18, 21, 26, 27, 31, 32, 53, 67), the factors “body site” and “age” in this data accounted for just 25.6% of total of the compositional variation between individuals within the flock. Multivariate regression model fitting with PERMANOVA indicated that age accounted for 20.3% of sample variance, while body site accounted for just 5.3%. “Flock” designation accounted for an additional 1.9% of variance. Studies in humans have also indicated that intrinsic factors such as age, sex, or body site account may be “significant” according to PERMANOVA while accounting for little overall variance (68, 69). In this instance, the large proportion of residual variance (72.6%) indicates influences from additional factors, such as natural variation between individual birds, variations in management, heterogeneity in the metacommunity around the birds in a field setting (70), and the introduction of artificial variation from the sampling methods used (44, 71, 72).

Nevertheless, this study has provided necessary data to set the groundwork for further studies that will enhance our understanding of turkey respiratory microbiome. Extensive metadata were collected throughout the study to track the flocks with regards to vaccination history, immune status, antibiotic treatment, weight gain trajectories, management, and so on. Despite our attempt to establish correlations or associations between microbiota and different metadata parameters, the uncontrolled field study has several limitations. Overlapping vaccination schedules impeded direct analysis of the effects of the vaccines on the microbiome, such as induction of inflammatory responses in the mucosal environment that may disrupt local microbial communities (73, 74). Likewise, we could not determine how microbiota responds to reovirus infection or other pathogens because the precise time of the natural infection was unknown, especially since the birds were also administered with live vaccines. Furthermore, the scope of this study did not accommodate assessment of different turkey breeds and production systems such as organic and pasture. In addition, although the QIIME 2 bioinformatics pipeline has allowed the clustering of 16S rRNA gene reads into biologically meaningful ASVs, curation of the reference databases (such as SILVA) used for taxonomic classification is still not rigorous enough to guarantee unequivocal taxonomic assignments. Likewise, the 16S rRNA gene sequencing and the sampling and sample processing methods employed in this study did not differentiate transient or dead occupants from live and active colonizers of the sampled sites. 16S rRNA barcoding methods in conjunction with microbial metabolomics profiles are one means of better phylotyping the resident microbiome (75).

## Conclusion

This study has presented a detailed picture of turkey respiratory microbiota over the course of a normal commercial turkey farm sequence. Body site and bird age together account for very little compositional variance between samples, suggesting that extrinsic factors (environmental and managerial) may influence beta diversity more substantially. Nevertheless, comparison of the composition and temporal dynamics of tracheal, nasal and gut microbiota in separate flocks has suggested several testable hypotheses: 1) that gut microbiota highly influence the occupation and pathogen susceptibility of respiratory mucosae via the surrounding environment; 2) that community structure in the nasal cavity is more stable than in the trachea, and therefore more easily manipulated; 3) that community assembly in the nasal cavity continuously influences what the trachea is exposed to; 4) that changes in immune status (via infection or vaccination) disrupt both the gut and respiratory microbiota, regardless of the site of infection. Additionally, by taking advantage of natural differences in the rates of flock weight gain trajectories, associations with poor flock performance were attributable to just a few taxa in both respiratory and gut sites. Future studies should focus on defining the effects of these taxa and elucidating the mechanisms of interfacing that may exist between the sites themselves, including influences from the environment.

## MATERIALS AND METHODS

### Flock management and sampling

Two flocks of commercial Hybrid Converter turkeys, each having more than 11,000 birds, were sampled for an entire production cycle between November 2015 and March 2016. The birds raised on this production cycle were all male. The birds were raised on a floor with wood shavings as litter and provided *ad libitum* access to feed and water. Feed formulations for each the brood and grow-out periods were not provided by the companyare proprietary information but are consistent between the two flocks. The flocks were reared in two adjacent barns (generically designated F1 and F2) during the brood period, i.e., from hatch to 5 weeks of age (WOA). After this period, each flock was moved to a different grow-out farm (Fig. 1). All farms are subsidiary to the same turkey production company, and therefore subject to the same management protocols and feeding regimens (including feed formulations). When referring to the flocks specifically during the brood stage, they are designated F1B and F2B. Likewise, when referring to the flocks during the grow-out stage, they are designated F1G and F2G, respectively.

Vaccinations during the brood period were performed on the same days for both flocks. The birds were vaccinated with AviPro® Megan® Egg (Elanco Animal Health Inc., Greenfield, IN, USA; against *Salmonella enterica* serovar Typhimurium) at 3 weeks, 4 days of age, and with H.E. Vac (Arko Labs, Ltd., Jewell, IA, USA; against hemorrhagic enteritis virus) at 4 WOA. During the grow-out period, the two farms administered vaccines and antimicrobials independently based on routine management protocol. Flock F1G was treated with AviPro® Megan® Egg at 9 WOA, Albendazole (Zoetis Inc., Parsippany, NJ, USA; for deworming) at 10 WOA, and two doses of M-NINEVAX®-C (Intervet Inc., Millsboro, DE, USA; against *Salmonella* and *Pasteurella multocida*) at 10 weeks, 2 days of age and again at 13 weeks, 6 days of age. Flock F2G was treated with chlortetracycline (CTC; Zoetis Inc., Parsippany, NJ, USA) at 7 weeks and 2 days, at 9 weeks and 4 days, and at 14 weeks and 2 days as a prophylactic measure against *Mycoplasma* infection. F2G was also given AviPro® Megan® Egg at 8 WOA and albendazole at 9 weeks, 4 days. Both grow-out flocks were administered with anticoccidial sulfadimethoxine (Di-Methox, Huvepharma, Inc., Peachtree City, GA, USA) in drinking water (Fig. 1).

A total of 104 birds were sampled at 3 time-points during brooding (1, 3, and 5 age WOA) and grow-out (8, 12, and 16 WOA). The sample sizes were as follows: 10 birds per flock at each time-point during brooding, 10 birds per flock at 8 WOA, 8 birds per flock at 12 WOA and 4 birds per flock at 16 WOA. Blood was drawn from live birds via the brachial (wing) vein for serum collection prior to euthanasia. The birds were humanely euthanized in accordance with protocol number 2015A00000056-R1 approved by The Ohio State University Institutional Animal Care and Use Committee. In brief, the animals were exposed to carbon dioxide (CO_2_) in a euthanasia chamber (1–3 birds, depending on bird size). The CO_2_ flow was maintained at 10– 30% displacement of chamber volume/minute until euthanasia was complete. Euthanasia was confirmed by absence of breathing and lack of heartbeat. After euthanasia, the body weight of each bird was measured separately prior to tissue excision.

### Detection of vaccine- and pathogen-specific serum antibodies

Serum samples were sent to the Animal Disease Diagnostic Lab, Ohio Department of Agriculture (Columbus, Ohio) for antibody quantification. Levels of reovirus antibodies in serum were measured by ELISA using IDEXX REO Ab Test kit (IDEXX, Westbrook, Maine, USA). Antibodies against the Newcastle disease virus (NDV), hemorrhagic enteritis virus (HEV), *Mycoplasma meleagridis* (MM), and *Mycoplasma synoviae* (MS) and *Mycoplasma gallisepticum* (MG) were quantified or detected with ProFLOK® NDV T Ab, HEV Ab, MM - T Ab, and MG-MS Ab kits (Zoetis), respectively.

### Sample collection, processing, DNA extraction, and 16Sr RNA gene sequencing

The ileum and cecum were each excised from the birds using sterile scissors and forceps. The tissues were placed in sterile Falcon tubes and immediately snap-frozen with liquid nitrogen. Deep-frozen cecum and ileum tissues were homogenized using mortar and pestle and subsampled to 0.3g total organic mass. DNA extraction for ileum and cecum homogenate (0.3g) were performed using the DNeasy PowerSoil Kit (Qiagen Sciences Inc., Germantown, MD).

The trachea was excised inside the biosafety cabinet and flushed by repeatedly drawing and expelling PBS through the length of the trachea (from both directions) with a filtered pipet tip. The nasal cavity was washed during necropsy by flushing and retrieving 150μL PBS through the nares with a 200μL filtered pipet tip. This process was repeated until about 1ml of total volume of PBS containing organic material was collected. Both tracheal and nasal wash collection solutions were centrifuged until a pellet of organic material formed at the bottom of the tube (10,000xg for 5 min). DNA extraction for tracheal and nasal wash pellets were performed using the DNeasy Blood & Tissue kit (Qiagen Sciences Inc., Germantown, MD). Samples of the same age and body site were extracted together, regardless of the flock designation.

DNA was quantified using NanoDrop 2000c Spectrophotometer (Thermo Fisher Scientific, Waltham, MA) and standardized to concentrations up to 100 ng/µL in a total volume of 20 µL/sample. 16S rRNA gene barcoding was performed with Illumina MiSeq (reagent kit v3; Illumina, San Diego, CA) by the University of Minnesota Genomics Center (Minneapolis, MN). Samples were not pooled prior to sequencing. Further details on collection, modifications to the default extraction protocols, PCR and sequencing conditions have been previously reported in Ngunjiri, et al. (2019) (27).

### Sequence processing and feature table generation and filtering

The barcoded Illumina sequencing data were demultiplexed to generate fastq files for each sample. Proximal and distal primers and adapters were trimmed from the reads using BBDuk, and paired-end reads were merged using BBMerge, both from the BBTools (v35) suite of DNA/RNA analysis software provided freely by the Joint Genome Institute (76). 16S rRNA amplicon primers were removed using Cutadapt (v1.4.2) (76, 77). Sequences of lengths less than 245 and greater than 260 base pairs were filtered using the BBMap module of BBTools.

The sequences were imported into QIIME2 (2019 v1.0) (78) in Casava 1.8 single-end demultiplexed format for further processing using different algorithms (implemented in QIIME2). Imported sequences were denoised with DADA2 (79) using the pooled-sample chimera filtering method. We used VSEARCH for detection of chimeric sequences and open-reference clustering of amplicon sequence variants (ASVs) with 100% identities using SILVA (release 132) reference database (80). The feature table produced from this step contained ASV counts for each sample. VSEARCH (81) was also used to identify non-16S rRNA gene ASVs, which were subsequently removed from the feature table. The table was filtered further to remove ASVs with a total frequency of <10 (in the entire table) and those present in only 1 sample. Next, the representative set of sequences were aligned *de novo* using MAFFT. Non-conserved and highly gapped columns of the alignment were removed by applying the “mast” option of the alignment plugin. The aligned filtered sequences were then used to reconstruct a phylogenetic tree using FastTree, which was rooted at the midpoint prior to application in beta diversity analysis.

For taxonomic classification, we trained a naïve Bayes classifier (82) on the 16S rRNA gene sequences spanning the region covered by the 515F (GTGYCAGCMGCCGCGGTAA) and 806R (GGACTACNVGGGTWTCTAAT) primer pair (83). The classifier was trained on reference sequences extracted from SILVA 16S rRNA gene database (release 132) based on matches to this primer pair and a taxonomic identification table (majority classification) based on reference sequences clustered at 99% identity. Lastly, the classifier was used to assign taxonomy to the ASVs observed in our experimental data.

The final feature table was rarefied to 5000 reads per sample. Samples with read counts under this amount were discarded. All further analyses were conducted in RStudio (84) with the following packages: BiocManager (85), biomformat (86), dplyr (87), reshape2 (88), ape (89), ggplot2 (90), gridExtra (91), RColorBrewer (92), ggpubr (93), xlsx (94), GUniFrac (95), vegan (96), VennDiagram (97), and pairwiseAdonis (98). Samples of the same body site, age group, and flock were considered a statistical population during analysis.

### Estimation of diversity and identification of predominant taxa in each body site

Alpha diversity indices taxonomic richness (number of ASVs per sample) and Pielou’s evenness were calculated using the ‘vegan’ package (99). Comparisons of value distributions between age groups were performed using a pairwise T-test with Bonferroni correction for multiple comparisons. The dependence of richness and evenness on age was assessed with linear regression. The response of the two flock groups was measured separately for each body site.

Comparisons between regressions from the same body site were performed using a modified T-test described by the following functions:

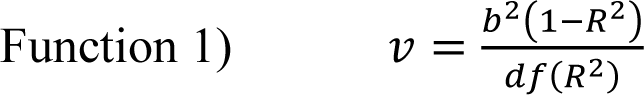

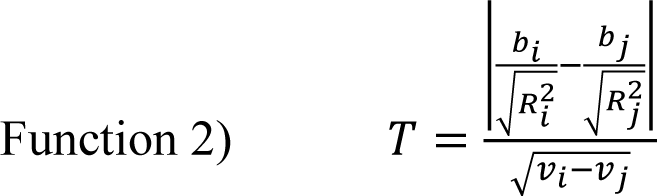

where the slope (*b*), degrees of freedom (*df*), and coefficient of determination (*R^2^*) for the regression line were used to calculate the variance (*v*), which was subsequently used to calculate Student’s *T*.

For beta diversity analysis, we used unweighted UniFrac distances, calculated by applying the midpoint rooted tree to the feature table (R package ‘GUniFrac’ (95)), to measure sample dispersion in composition space. Dimensionality was reduced using principal coordinates analysis (PCO; R package ‘vegan’ (96)), and differences between sample groups were quantified with pairwise Permutational Multivariate Analysis of Variance (PERMANOVA; R package ‘pairwiseAdonis’ (98)).

The distribution of all ASVs among body sites was visualized using a four-set Venn diagram (R package ‘VennDiagram’ (97)). Selection of predominant taxa was performed independently for each body site, using a serial filtering scheme. The first filter pulled ASVs that were present in more than 50% of samples within an age group. The second filter retained ASVs that were common among the filtered lists of all age groups. The third filter retained only those ASVs that were present at a mean relative abundance of more than 3.5% in at least one age group. This process was performed for several selections of consecutive age groups.

### Data availability statement

Raw data files and metadata are publicly available in the NCBI BioProject database under accession number PRJNA589216 at https://www.ncbi.nlm.nih.gov/bioproject/589216. Raw data files are also available through Sequence Read Archive (SRA) under accession numbers SAMN13269218 through SAMN13269601. Pipe scripts for QIIME2 sequence processing and generation of ASV table and R scripts for analysis are publicly available under at https://github.com/kjmtaylor22/cwl-cats/tree/master.

## SUPPLEMENTAL MATERIAL

Supplemental material for this article may be found at

**Supplemental file 1**, PDF file, 11.4 MB.

**Supplemental file 2**, PDF file, 328 KB.

## ACKNOWLEDGEMENTS

We thank the management team of the commercial farm participating in this study for providing turkeys and giving us access to flock treatment and management data related to this study. The identity of the commercial farm referenced in this study will remain anonymous as a condition of their open sharing of rearing metadata and cooperation during sampling.

All animals used in this study were collected with consent from the commercial farm and were humanely euthanized in accordance with protocol number 2015A00000056-R1 approved by The Ohio State University Institutional Animal Care and Use Committee. We declare no conflicts of interest.

Sequence processing and analysis were performed using the resources of the Molecular & Cellular Imaging Center, Ohio Agricultural Research and Development Center, The Ohio State University.

This project was supported by Agriculture and Food Research Initiative competitive grant number 2015-68004-23131 from the USDA National Institute of Food and Agriculture (to C.-W.L. and T.J.J.). C.-W.L. and T.J.J. developed the broad concept for the study. C.-W.L., J.M.N., and M.C.A. designed and managed the study. J.M.N., M.C.A., H.J., M.E., M.K.C., and C.-W.L. collected the samples. J.M.N., M.C.A., and A.G. performed sample processing and DNA extraction. B.P.W. and T.J.J. managed the DNA sequencing. K.J.M.T. and J.M.N. performed bioinformatics analyses. K.J.M.T. and J.M.N. wrote the manuscript with input from all authors. C.-W.L. provided reagents and sampling resources. All authors read and approved the final manuscript.

## REFERENCES

1. Jankowski J, Zdunczyk Z, Mikulski D, Przybylska-Gornowicz B, Sosnowska E, Juskiewicz J. 2013. Effect of whole wheat feeding on gastrointestinal tract development and performance of growing turkeys. Anim Feed Sci Technol 185:150–159.

2. Lei K, Li YL, Yu DY, Rajput IR, Li WF. 2013. Influence of dietary inclusion of Bacillus licheniformis on laying performance, egg quality, antioxidant enzyme activities, and intestinal barrier function of laying hens. Poult Sci 92:2389–2395.

3. Celi P, Cowieson AJ, Fru-Nji F, Steinert RE, Kluenter A-M, Verlhac V. 2017. Gastrointestinal functionality in animal nutrition and health: New opportunities for sustainable animal production. Animal Feed Science and Technology 234:88–100.

4. Choi KY, Lee TK, Sul WJ. 2015. Metagenomic Analysis of Chicken Gut Microbiota for Improving Metabolism and Health of Chickens — A Review. Asian-Australas J Anim Sci 28:1217–1225.

5. Simon K, Verwoolde MB, Zhang J, Smidt H, de Vries Reilingh G, Kemp B, Lammers A. 2016. Long-term effects of early life microbiota disturbance on adaptive immunity in laying hens. Poultry Science 95:1543–1554.

6. Danzeisen JL, Calvert AJ, Noll SL, McComb B, Sherwood JS, Logue CM, Johnson TJ. 2013. Succession of the turkey gastrointestinal bacterial microbiome related to weight gain. PeerJ 1:e237.

7. Kurtoglu V, Kurtoglu F, Seker E, Coskun B, Balevi T, Polat ES. 2004. Effect of probiotic supplementation on laying hen diets on yield performance and serum and egg yolk cholesterol. Food Additives & Contaminants 21:817–823.

8. Pan D, Yu Z. 2014. Intestinal microbiome of poultry and its interaction with host and diet. Gut Microbes 5:108–119.

9. Borda-Molina D, Seifert J, Camarinha-Silva A. 2018. Current Perspectives of the Chicken Gastrointestinal Tract and Its Microbiome. Computational and Structural Biotechnology Journal 16:131–139.

10. Oakley BB, Lillehoj HS, Kogut MH, Kim WK, Maurer JJ, Pedroso A, Lee MD, Collett SR, Johnson TJ, Cox NA. 2014. The chicken gastrointestinal microbiome. FEMS Microbiol Lett 360:100–112.

11. Scupham AJ. 2009. Campylobacter Colonization of the Turkey Intestine in the Context of Microbial Community Development. Appl Environ Microbiol 75:3564–3571.

12. Scupham AJ. 2007. Succession in the intestinal microbiota of preadolescent turkeys. FEMS Microbiol Ecol 60:136–147.

13. D’Andreano S, Sànchez Bonastre A, Francino O, Cuscó Martí A, Lecchi C, Grilli G, Giovanardi D, Ceciliani F. 2017. Gastrointestinal microbial population of turkey (Meleagris gallopavo) affected by hemorrhagic enteritis virus. Poultry Science 96:3550–3558.

14. Calvert AJ. 2012. Light turkey syndrome: field study and inoculation trial.

15. Zdunczyk Z, Jankowski J, Kaczmarek S, Juskiewicz J. 2015. Determinants and effects of postileal fermentation in broilers and turkeys part 1: gut microbiota composition and its modulation by feed additives. Worlds Poult Sci J 71:37–47.

16. Ward TL, Weber BP, Mendoza KM, Danzeisen JL, Llop K, Lang K, Clayton JB, Grace E, Brannon J, Radovic I, Beauclaire M, Heisel TJ, Knights D, Cardona C, Kogut M, Johnson C, Noll SL, Arsenault R, Reed KM, Johnson TJ. 2019. Antibiotics and Host-Tailored Probiotics Similarly Modulate Effects on the Developing Avian Microbiome, Mycobiome, and Host Gene Expression. mBio 10:e02171–19.

17. Danzeisen JL, Clayton JB, Huang H, Knights D, McComb B, Hayer SS, Johnson TJ. 2015. Temporal Relationships Exist Between Cecum, Ileum, and Litter Bacterial Microbiomes in a Commercial Turkey Flock, and Subtherapeutic Penicillin Treatment Impacts Ileum Bacterial Community Establishment. Frontiers in Veterinary Science 2.

18. Wilkinson TJ, Cowan AA, Vallin HE, Onime LA, Oyama LB, Cameron SJ, Gonot C, Moorby JM, Waddams K, Theobald VJ, Leemans D, Bowra S, Nixey C, Huws SA. 2017. Characterization of the Microbiome along the Gastrointestinal Tract of Growing Turkeys. Frontiers in Microbiology 8.

19. Niedermeyer JA, Ring L, Miller WG, Genger S, Lindsey CP, Osborne J, Kathariou S. 2018. Proximity to Other Commercial Turkey Farms Affects Colonization Onset, Genotypes, and Antimicrobial Resistance Profiles of Campylobacter spp. in Turkeys: Suggestive Evidence from a Paired-Farm Model. Applied and Environmental Microbiology 84.

20. Smith AH, Rehberger TG. 2018. Bacteria and fungi in day-old turkeys vary among companies, collection periods, and breeder flocks. Poult Sci 97:1400–1411.

21. Donaldson EE, Stanley D, Hughes RJ, Moore RJ. 2017. The time-course of broiler intestinal microbiota development after administration of cecal contents to incubating eggs. PeerJ 5:e3587.

22. Hieke A-SC, Hubert SM, Athrey G. 2019. Circadian disruption and divergent microbiota acquisition under extended photoperiod regimens in chicken. PeerJ 7:e6592.

23. Wang L, Lilburn M, Yu Z. 2016. Intestinal Microbiota of Broiler Chickens As Affected by Litter Management Regimens. Frontiers in Microbiology 7.

24. Cressman MD, Yu Z, Nelson MC, Moeller SJ, Lilburn MS, Zerby HN. 2010. Interrelations between the Microbiotas in the Litter and in the Intestines of Commercial Broiler Chickens. Applied and Environmental Microbiology 76:6572–6582.

25. Danzeisen JL, Kim HB, Isaacson RE, Tu ZJ, Johnson TJ. 2011. Modulations of the Chicken Cecal Microbiome and Metagenome in Response to Anticoccidial and Growth Promoter Treatment. PLOS ONE 6:e27949.

26. Johnson TJ, Youmans BP, Noll S, Cardona C, Evans NP, Karnezos TP, Ngunjiri JM, Abundo MC, Lee C-W. 2018. A Consistent and Predictable Commercial Broiler Chicken Bacterial Microbiota in Antibiotic-Free Production Displays Strong Correlations with Performance. Applied and Environmental Microbiology 84.

27. Ngunjiri JM, Taylor KJM, Abundo MC, Jang H, Elaish M, Kc M, Ghorbani A, Wijeratne S, Weber BP, Johnson TJ, Lee C-W. 2019. Farm Stage, Bird Age, and Body Site Dominantly Affect the Quantity, Taxonomic Composition, and Dynamics of Respiratory and Gut Microbiota of Commercial Layer Chickens. Appl Environ Microbiol 85:e03137–18.

28. Naylor CJ, Al-Ankari AR, Al-Afaleq AI, Bradbury JM, Jones RC. 1992. Exacerbation of Mycoplasma gallisepticum infection in Turkeys by rhinotracheitis virus. Avian Pathology 21:295–305.

29. Power J, Jordan FTW. 1976. A comparison of the virulence of three strains of Mycoplasma gallisepticum and one strain of Mycoplasma gallinarum in chicks, turkey poults, tracheal organ cultures and embryonated fowl eggs. Research in Veterinary Science 21:41–46.

30. De Rosa M, Droual R, Chin RP, Shivaprasad HL, Walker RL. 1996. Ornithobacterium rhinotracheale Infection in Turkey Breeders. Avian Diseases 40:865–874.

31. Glendinning L, McLachlan G, Vervelde L. 2017. Age-related differences in the respiratory microbiota of chickens. PLOS ONE 12:e0188455.

32. Shabbir MZ, Malys T, Ivanov YV, Park J, Shabbir MAB, Rabbani M, Yaqub T, Harvill ET. 2015. Microbial communities present in the lower respiratory tract of clinically healthy birds in Pakistan. Poult Sci 94:612–620.

33. Kallapura G, Hernandez-Velasco X, Pumford N, Bielke L, Hargis BM, Tellez G. 2014. Evaluation of respiratory route as a viable portal of entry for Salmonella in poultry. Vet Med Res Rep 5:59–79.

34. Pantin-Jackwood MJ, Day JM, Jackwood MW, Spackman E. 2008. Enteric Viruses Detected by Molecular Methods in Commercial Chicken and Turkey Flocks in the United States Between 2005 and 2006. Avian Diseases 52:235–244.

35. Ferguson-Noel N. 2013. Mycoplasmosis, p. 875–941. In Diseases of poultry. John Wiley & Sons, Ames, IA.

36. Hybrid Turkeys. 2019. Performance Goals: Converter Commercial Males.

37. Yavari CA, Ramírez AS, Nicholas RAJ, Radford AD, Darby AC, Bradbury JM. 2017. Mycoplasma tullyi sp. nov., isolated from penguins of the genus Spheniscus. International Journal of Systematic and Evolutionary Microbiology, 67:3692–3698.

38. Bradbury JM, Abdul-Wahab OMS, Yavari CA, Dupiellet J-P, Bové JM. 1993. Mycoplasma imitans sp. nov. Is Related to Mycoplasma gallisepticum and Found in Birds. International Journal of Systematic and Evolutionary Microbiology, 43:721–728.

39. Abdul-Wahab OM, Ross G, Bradbury JM. 1996. Pathogenicity and cytadherence of Mycoplasma imitans in chicken and duck embryo tracheal organ cultures. Infection and Immunity 64:563–568.

40. Glisson. 1998. Bacterial respiratory disease of poultry. Poult Sci 77:1139–1142.

41. Blackall PJ, Soriano-Vargas EV. 2013. Infectious coryza and related bacterial infections, p. 859–873. In Diseases of Poultry. John Wiley & Sons, Ames, IA.

42. van Empel P, van den Bosch H, Goovaerts D, Storm P. 1996. Experimental infection in turkeys and chickens with Ornithobacterium rhinotracheale. Avian Dis 40:858–864.

43. Rosenstein R, Götz F. 2013. What distinguishes highly pathogenic staphylococci from medium- and non-pathogenic? Curr Top Microbiol Immunol 358:33–89.

44. Salter SJ, Cox MJ, Turek EM, Calus ST, Cookson WO, Moffatt MF, Turner P, Parkhill J, Loman NJ, Walker AW. 2014. Reagent and laboratory contamination can critically impact sequence-based microbiome analyses. BMC Biology 12:87.

45. Liang R, Xiao P, She R, Han S, Chang L, Zheng L. 2013. Culturable Airborne Bacteria in Outdoor Poultry-Slaughtering Facility. Microbes and Environments advpub.

46. Luyckx K, Van Coillie E, Dewulf J, Van Weyenberg S, Herman L, Zoons J, Vervaet E, Heyndrickx M, De Reu K. 2017. Identification and biocide susceptibility of dominant bacteria after cleaning and disinfection of broiler houses. Poultry Science 96:938–949.

47. Sghaier H, Bouchami O, Desler C, Lazim H, Saidi M, Rasmussen LJ, Ben Hassen A. 2012. Analysis of the antimicrobial susceptibility of the ionizing radiation-resistant bacterium Deinococcus radiodurans: implications for bioremediation of radioactive waste. Ann Microbiol 62:493–500.

48. Angelakis E, Raoult D. 2010. The Increase of Lactobacillus Species in the Gut Flora of Newborn Broiler Chicks and Ducks Is Associated with Weight Gain. PLOS ONE 5:e10463.

49. Torok VA, Ophel-Keller K, Loo M, Hughes RJ. 2008. Application of Methods for Identifying Broiler Chicken Gut Bacterial Species Linked with Increased Energy Metabolism. Appl Environ Microbiol 74:783–791.

50. Torok VA, Hughes RJ, Mikkelsen LL, Perez-Maldonado R, Balding K, MacAlpine R, Percy NJ, Ophel-Keller K. 2011. Identification and Characterization of Potential Performance-Related Gut Microbiotas in Broiler Chickens across Various Feeding Trials. Applied and Environmental Microbiology 77:5868–5878.

51. Thomason CA, Mullen N, Belden LK, May M, Hawley DM. 2017. Resident Microbiome Disruption with Antibiotics Enhances Virulence of a Colonizing Pathogen. Sci Rep 7:1–8.

52. Chang CLT, Chung C-Y, Kuo C-H, Kuo T-F, Yang C-W, Yang W-C. 2016. Beneficial Effect of Bidens pilosa on Body Weight Gain, Food Conversion Ratio, Gut Bacteria and Coccidiosis in Chickens. PLOS ONE 11:e0146141.

53. Lu J, Idris U, Harmon B, Hofacre C, Maurer JJ, Lee MD. 2003. Diversity and Succession of the Intestinal Bacterial Community of the Maturing Broiler Chicken. Applied and Environmental Microbiology 69:6816–6824.

54. Martynova-Van Kley MA, Oviedo-Ron EO, Dowd SE, Hume M, Nalian A. 2012. Effect of Eimeria Infection on Cecal Microbiome of Broilers Fed Essential Oils. International Journal of Poultry Science 11:747–755.

55. Allen-Vercoe E, Strauss J, Chadee K. 2011. Fusobacterium nucleatum. Gut Microbes 2:294–298.

56. Chen Y-J, Wu H, Wu S-D, Lu N, Wang Y-T, Liu H-N, Dong L, Liu T-T, Shen X-Z. 2018. Parasutterella, in association with irritable bowel syndrome and intestinal chronic inflammation. Journal of Gastroenterology and Hepatology 33:1844–1852.

57. Langworth BF. 1977. Fusobacterium necrophorum: its characteristics and role as an animal pathogen. Bacteriol Rev 41:373–390.

58. Strauss J, Kaplan GG, Beck PL, Rioux K, Panaccione R, DeVinney R, Lynch T, Allen-Vercoe E. 2011. Invasive potential of gut mucosa-derived fusobacterium nucleatum positively correlates with IBD status of the host. Inflamm Bowel Dis 17:1971–1978.

59. Costello EK, Stagaman K, Dethlefsen L, Bohannan BJM, Relman DA. 2012. The Application of Ecological Theory Toward an Understanding of the Human Microbiome. Science 336:1255–1262.

60. Huxley EJ, Viroslav J, Gray WR, Pierce AK. 1978. Pharyngeal aspiration in normal adults and patients with depressed consciousness. The American Journal of Medicine 64:564–568.

61. Venkataraman A, Bassis CM, Beck JM, Young VB, Curtis JL, Huffnagle GB, Schmidt TM. 2015. Application of a Neutral Community Model To Assess Structuring of the Human Lung Microbiome. mBio 6:e02284–14.

62. Dickson RP, Erb-Downward JR, Martinez FJ, Huffnagle GB. 2016. The Microbiome and the Respiratory Tract. Annual Review of Physiology 78:481–504.

63. Yatsunenko T, Rey FE, Manary MJ, Trehan I, Dominguez-Bello MG, Contreras M, Magris M, Hidalgo G, Baldassano RN, Anokhin AP, Heath AC, Warner B, Reeder J, Kuczynski J, Caporaso JG, Lozupone CA, Lauber C, Clemente JC, Knights D, Knight R, Gordon JI. 2012. Human gut microbiome viewed across age and geography. Nature 486:222–227.

64. Awad WA, Mann E, Dzieciol M, Hess C, Schmitz-Esser S, Wagner M, Hess M. 2016. Age-Related Differences in the Luminal and Mucosa-Associated Gut Microbiome of Broiler Chickens and Shifts Associated with Campylobacter jejuni Infection. Front Cell Infect Microbiol 6.

65. Odamaki T, Kato K, Sugahara H, Hashikura N, Takahashi S, Xiao J, Abe F, Osawa R. 2016. Age-related changes in gut microbiota composition from newborn to centenarian: a cross-sectional study. BMC Microbiology 16:90.

66. Dumas MD, Polson SW, Ritter D, Ravel J, Gelb J, Morgan R, Wommack KE. 2011. Impacts of Poultry House Environment on Poultry Litter Bacterial Community Composition. PLoS ONE 6:e24785.

67. Oakley BB, Buhr RJ, Ritz CW, Kiepper BH, Berrang ME, Seal BS, Cox NA. 2014. Successional changes in the chicken cecal microbiome during 42 days of growth are independent of organic acid feed additives. BMC Veterinary Research 10:282.

68. Zhang X, Zhong H, Li Y, Shi Z, Zhang Z, Zhou X, others. Age-dependent sexual dimorphism in the adult human gut microbiota. BioRxiv 2019: 646620.

69. Mahnic A, Rupnik M. 2018. Different host factors are associated with patterns in bacterial and fungal gut microbiota in Slovenian healthy cohort. PLOS ONE 13:e0209209.

70. Miller ET, Svanback R, Bohannan BJM. 2018. Microbiomes as Metacommunities: Understanding Host-Associated Microbes through Metacommunity Ecology. Trends Ecol Evol 33:926–935.

71. Callahan BJ, McMurdie PJ, Rosen MJ, Han AW, Johnson AJA, Holmes SP. 2016. DADA2: High resolution sample inference from Illumina amplicon data. Nat Methods 13:581–583.

72. Ranjan R, Rani A, Metwally A, McGee HS, Perkins DL. 2016. Analysis of the microbiome: Advantages of whole genome shotgun versus 16S amplicon sequencing. Biochem Biophys Res Commun 469:967–977.

73. Park SH, Kim SA, Rubinelli PM, Roto SM, Ricke SC. 2017. Microbial compositional changes in broiler chicken cecal contents from birds challenged with different Salmonella vaccine candidate strains. Vaccine 35:3204–3208.

74. Tarabichi Y, Li K, Hu S, Nguyen C, Wang X, Elashoff D, Saira K, Frank B, Bihan M, Ghedin E, Methé BA, Deng JC. 2015. The administration of intranasal live attenuated influenza vaccine induces changes in the nasal microbiota and nasal epithelium gene expression profiles. Microbiome 3:74.

75. Turnbaugh PJ, Gordon JI. 2008. An Invitation to the Marriage of Metagenomics and Metabolomics. Cell 134:708–713.

76. Bushnell B, Rood J, Singer E. 2017. BBMerge – Accurate paired shotgun read merging via overlap. PLOS ONE 12:e0185056.

77. Martin M. 2011. Cutadapt removes adapter sequences from high-throughput sequencing reads. EMBnet.journal 17:10.

78. Bolyen E, Rideout JR, Dillon MR, Bokulich NA, Abnet CC, Al-Ghalith GA, Alexander H, Alm EJ, Arumugam M, Asnicar F, Bai Y, Bisanz JE, Bittinger K, Brejnrod A, Brislawn CJ, Brown CT, Callahan BJ, Caraballo-Rodríguez AM, Chase J, Cope EK, Da Silva R, Diener C, Dorrestein PC, Douglas GM, Durall DM, Duvallet C, Edwardson CF, Ernst M, Estaki M, Fouquier J, Gauglitz JM, Gibbons SM, Gibson DL, Gonzalez A, Gorlick K, Guo J, Hillmann B, Holmes S, Holste H, Huttenhower C, Huttley GA, Janssen S, Jarmusch AK, Jiang L, Kaehler BD, Kang KB, Keefe CR, Keim P, Kelley ST, Knights D, Koester I, Kosciolek T, Kreps J, Langille MGI, Lee J, Ley R, Liu Y-X, Loftfield E, Lozupone C, Maher M, Marotz C, Martin BD, McDonald D, McIver LJ, Melnik AV, Metcalf JL, Morgan SC, Morton JT, Naimey AT, Navas-Molina JA, Nothias LF, Orchanian SB, Pearson T, Peoples SL, Petras D, Preuss ML, Pruesse E, Rasmussen LB, Rivers A, Robeson MS, Rosenthal P, Segata N, Shaffer M, Shiffer A, Sinha R, Song SJ, Spear JR, Swafford AD, Thompson LR, Torres PJ, Trinh P, Tripathi A, Turnbaugh PJ, Ul-Hasan S, van der Hooft JJJ, Vargas F, Vázquez-Baeza Y, Vogtmann E, von Hippel M, Walters W, Wan Y, Wang M, Warren J, Weber KC, Williamson CHD, Willis AD, Xu ZZ, Zaneveld JR, Zhang Y, Zhu Q, Knight R, Caporaso JG. 2019. Reproducible, interactive, scalable and extensible microbiome data science using QIIME 2. Nat Biotechnol 37:852–857.

79. Callahan BJ, McMurdie PJ, Rosen MJ, Han AW, Johnson AJA, Holmes SP. 2016. DADA2: High-resolution sample inference from Illumina amplicon data. Nature Methods 13:581–583.

80. Quast C, Pruesse E, Yilmaz P, Gerken J, Schweer T, Yarza P, Peplies J, Glöckner FO. 2012. The SILVA ribosomal RNA gene database project: improved data processing and web-based tools. Nucleic Acids Research 41:D590–D596.

81. Rognes T, Flouri T, Nichols B, Quince C, Mahé F. 2016. VSEARCH: a versatile open source tool for metagenomics. PeerJ 4:e2584.

82. Bokulich NA, Kaehler BD, Rideout JR, Dillon M, Bolyen E, Knight R, Huttley GA, Caporaso JG. 2018. Optimizing taxonomic classification of marker-gene amplicon sequences with QIIME 2’s q2-feature-classifier plugin. Microbiome 6.

83. Caporaso JG, Lauber CL, Walters WA, Berg-Lyons D, Huntley J, Fierer N, Owens SM, Betley J, Fraser L, Bauer M, Gormley N, Gilbert JA, Smith G, Knight R. 2012. Ultra-high-throughput microbial community analysis on the Illumina HiSeq and MiSeq platforms. The ISME Journal 6:1621–1624.

84. R Core Team. 2019. R: A Language and Environment for Statistical Computing. R Foundation for Statistical Computing, Vienna, Austria.

85. Morgan M. 2018. BiocManager: Access the Bioconductor Project Package Repository.

86. McMurdie PJ, Paulson JN. 2019. biomformat: An interface package for the BIOM file format.

87. Wickham H, Francois R, Henry L, Muller K. 2019. dplyr: A Grammar of Data Manipulation.

88. Wickham H. 2007. Reshaping Data with the reshape Package. Journal of Statistical Software 21:1–20.

89. Paradis E, Schliep K. 2018. ape 5.0: an environment for modern phylogenetics and evolutionary analyses in R. Bioinformatics 35:526–528.

90. Wickham H. 2016. ggplot2: Elegant Graphics for Data Analysis. Springer-Verlag New York.

91. Auguie B. 2017. gridExtra: Miscellaneous Functions for “Grid” Graphics.

92. Neuwirth E. 2014. RColorBrewer: ColorBrewer Palettes.

93. Kassambara A. 2019. ggpubr: “ggplot2” Based Publication Ready Plots.

94. Dragulescu AA, Arendt C. 2018. xlsx: Read, Write, Format Excel 2007 and Excel 97/2000/XP/2003 Files.

95. Chen J. 2018. GUniFrac: Generalized UniFrac Distances.

96. Oksanen J, Blanchet FG, Friendly M, Kindt R, Legendre P, McGlinn D, Minchin PR, O’Hara RB, Simpson GL, Solymos P, Stevens MHH, Szoecs E, Wagner H. 2019. vegan: Community Ecology Package.

97. Chen H. 2018. VennDiagram: Generate High-Resolution Venn and Euler Plots.

98. Martinez Arbizu P. 2017. pairwiseAdonis: Pairwise Multilevel Comparison using Adonis.

99. Oksanen J, Blanchet FG, Friendly M, Kindt R, Legendre P, McGlinn D, Minchin PR, O’Hara RB, Simpson GL, Solymos P, Stevens MHH, Szoecs E, Wagner H. 2019. vegan: Community Ecology Package.

